# A comprehensive mass spectral library for human thyroid tissues

**DOI:** 10.1101/2021.05.27.445992

**Authors:** Yaoting Sun, Lu Li, Weigang Ge, Zhen Dong, Wei Liu, Hao Chen, Qi Xiao, Xue Cai, Fangfei Zhang, Junhong Xiao, Guangzhi Wang, Yi He, Oi Lian Kon, N. Gopalakrishna Iyer, Yongfu Zhao, Tiannan Guo

**Affiliations:** Zhejiang University, Hangzhou, China; Key Laboratory of Structural Biology of Zhejiang Province, School of Life Sciences, Westlake University, Hangzhou 310024, China; Center for Infectious Disease Research, Westlake Laboratory of Life Sciences and Biomedicine, Hangzhou 310024, China; Institute of Basic Medical Sciences, Westlake Institute for Advanced Study, Hangzhou 310024, China; Westlake Omics (Hangzhou) Biotechnology Co., Ltd., Hangzhou 310024, China; Department of General Surgery, The Second Hospital of Dalian Medical University, Dalian, China; Department of Urology, The Second Hospital of Dalian Medical University, Dalian, China; Division of Medical Sciences, National Cancer Centre Singapore, Republic of Singapore; Department of Head and Neck Surgery, National Cancer Centre Singapore, Republic of Singapore

## Abstract

Thyroid nodules occur in about 60% of the population. Current diagnostic strategies, however, often fail at distinguishing malignant nodules before surgery, thus leading to unnecessary, invasive treatments. As proteins are involved in all physio/pathological processes, a proteome investigation of biopsied nodules may help correctly classify and identify malignant nodules and discover therapeutic targets. Quantitative mass spectrometry data-independent acquisition (DIA) enables highly reproducible and rapid throughput investigation of proteomes. An exhaustive spectral library of thyroid nodules is essential for DIA yet still unavailable. This study presents a comprehensive thyroid spectral library covering five types of thyroid tissue: multinodular goiter, follicular adenoma, follicular and papillary thyroid carcinoma, and normal thyroid tissue. Our library includes 925,330 transition groups, 157,548 peptide precursors, 121,960 peptides, 9941 protein groups, and 9826 proteins from proteotypic peptides. This library resource was evaluated using three papillary thyroid carcinoma samples and their corresponding adjacent normal thyroid tissue, leading to effective quantification of up to 7863 proteins from biopsy-level thyroid tissues.

## Background & Summary

Thyroid nodules are common and, given the sensitivity of current diagnostic techniques, can be detected in approximately 60% of the general population, especially in women^1,2^. The incidence of thyroid malignancy or thyroid carcinoma, has rapidly increased over the last decades, although it is uncertain if this is a real increase or simply a result of widespread use of screening ultrasonography^3,4^. Most of these nodules are asymptomatic. Only 4-7% of patients present with complaints attributed to thyroid nodules. Although ultrasonography and ultrasound-guided fine-needle aspiration can help distinguish between benign and malignant nodules, approximately 30% of thyroid nodules remain indeterminate by cytopathology and require diagnostic surgery^5^, after which histopathology of surgical specimens provides a definitive and complete diagnosis. More importantly, only 15% of indeterminate nodules prove to be malignant. Because many benign nodules are clinically ambiguous and a source of uncertainty, such patients often undergo unnecessary surgery. Nucleic acid-based molecular tests, which require next-generation sequencing technology^6^, are currently used in clinical practice to reduce overtreatment of thyroid nodules. However, the diagnostic specificity of these tests remains modest at best (40-70%) for a myriad of reasons.

Unlike nucleic acids, proteins are directly involved in all life processes and determine cellular and organismal phenotype. Proteins also have the potential to be critical biomarkers for disease diagnosis and are themselves potential drug targets. For these reasons, there is tremendous potential in exploring thyroid molecular pathology from a protein-based perspective. Mass spectrometry (MS) -based proteomics has reached a high level of technical and methodological development during the last decade. Data-independent acquisition (DIA), in particular, enables comprehensive quantitation of peptides from complex compositions with high reproducibility and throughput^7^. In the conventional data-dependent acquisition (DDA) mode, only peptide precursors with high abundance in MS1 are fragmented. In DIA, however, all precursors within a predefined range (also called window) of mass-to-charge ratio (*m/z*) are fragmented by sequentially repeated cycling in windows, thus providing detailed data without loss of any eluted peptides^7,8^. Our group’s established pressure cycling technology (PCT)-based sample preparation methodology, coupled with DIA-MS, achieves the acquisition of complete proteomic information in less than six hrs^9,10^.

To optimize the efficiency of spectral identifications, DIA data analysis requires tissue- or organism-specific spectral libraries^8,11^. Although a pan-human library derived from healthy subjects has already been established^12,13^, this extensive and non-specific library could cause inaccuracies during ion matching. In recent years, several novel software for DIA data analysis, such as DIA-Umpire^14^, PECAN^15^, or DIA-NN^16^, no longer require spectral libraries. However, this library-free mode should be applied with caution due to its lower sensitivity and protein identification power compared to a library-based strategy^17^. A tissue-specific library for thyroid nodules, both benign and malignant, as well as for healthy thyroid, would thus provide an essential resource for the proteomic investigation of thyroid pathologies in a high-throughput manner.

This study introduces a thyroid-specific spectral library to support protein identification and quantification in thyroid nodules by DIA-MS (Figure 1). Five types of thyroid tissues were collected, namely normal tissue, two types of benign nodules (multinodular goiter (MNG) and follicular adenoma (FA), and two types of thyroid carcinomas (follicular thyroid carcinoma (FTC) and papillary thyroid carcinoma (PTC). Normal thyroid and thyroid nodule tissues were processed by PCT; extracted and desalted peptides were then combined into three different pooled samples: (1) pooled sample containing all five types, (2) PTC pooled samples, and (3) FA and FTC pooled sample. The pooled peptides were fractionated in two ways, *i*.*e*. strong cation exchange (SCX) or high-pH reversed-phase chromatography, to achieve higher peptide coverage. Peptide fractions were injected into HPLC-MS/MS with 60 min-gradient through DDA mode using Thermo Orbitrap Q Exactive™ HF. 46 DDA files were acquired in total. Our spectral library was built with Spectronaut 14.6 and included 925,330 transition groups, 157,548 precursors, 121,960 peptides, 9941 protein groups, and 9826 proteins from proteotypic peptides. We then validated this library by applying it to four DIA datasets acquired with four different methods (Figure 1).

**Figure 1.**
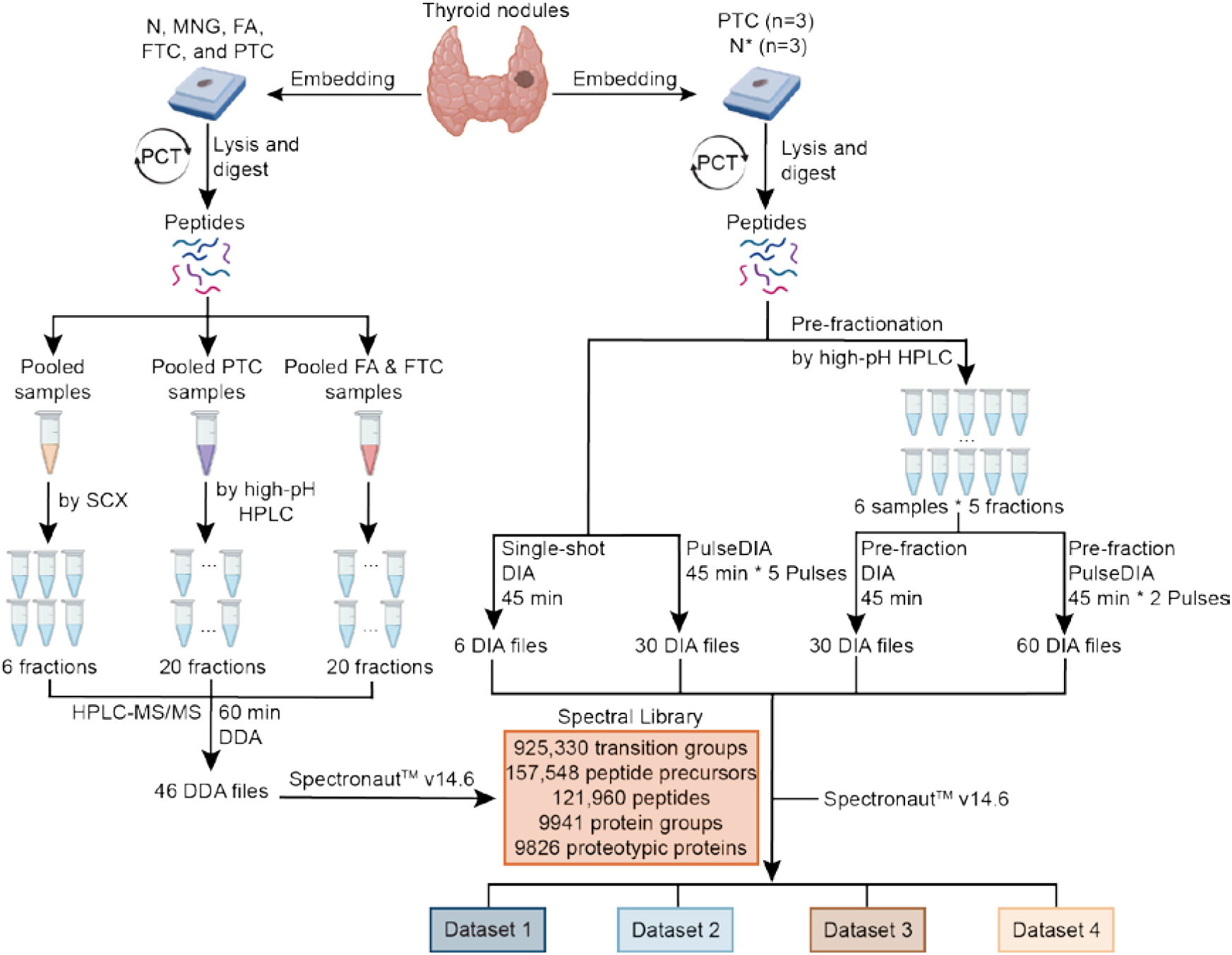
Workflow for generating a comprehensive thyroid-specific spectral library (left) and for its validation (right).

## Methods

### Sample collection

For the spectral library construction and testing, thyroid healthy and nodular samples were collected, between 2011 and 2019, from two clinical centers in Singapore (Singapore General Hospital) and China (The Second Hospital of Dalian Medical University). Ethical approval was given by both hospitals. Tissue cores of 1 mm diameter (0.6-1.2 mg) were extracted from the pathological regions of interest in formalin-fixed paraffin-embedded (FFPE) tissue blocks demarcated by experienced histopathologists^18^. Four types of thyroid nodules (42 MNG, 49 FA, 33 FTC, and 54 PTC) and 10 normal thyroid tissues were used for building the library. We also collected three paired PTC and corresponding tumor-adjacent tissues for validation of the spectral library.

### Sample preparation assisted by PCT

Samples were dewaxed, hydrated, and acidified using, in sequence, heptane, a decreasing ethanol series (100%, 90%, and 75%), and formic acid. The samples were next kept under basic hydrolysis conditions in Tris-HCl (100 mM, pH = 10) at 95 °C for 30 min, then transferred into a solution containing 30 µL lysis buffer (6 M urea, 2 M thiourea), 5 µL tris(2-carboxyethyl)phosphine (TECP, 10 mM), and 2.5 µL iodoacetamide (IAA) (40 mM). In PCT-Micro tubes, samples were lysed, reduced, and hydroxylated at 30 °C using PCT (120 cycles, 45 Kpsi, 30 s on-time, 10 s off-time). Trypsin (enzyme-to-substrate ratio, 1:50; Hualishi Scientific, China) and LysC (enzyme-to-substrate ratio, 1:40; Hualishi Scientific, China) were then added, followed by PCT-assisted digestion (120 cycles, 20 Kpsi, 50 s on-time, 10 s off-time). 1% trifluoroacetic acid (TFA) was added to terminate the digestion process. The resulting peptides were desalted with 2% acetonitrile (ACN) and 0.1% TFA and reconstituted with 2% ACN containing 0.1% formic acid. Peptide concentrations were measured by Nanoscan (Analytic Jena, Germany) at A_280_, and samples were stored at 4 °C for further analysis. For sample testing, we used previously optimized methods^10,19^. All the chemical reagents were obtained from Sigma-Aldrich.

### Strong cation exchange (SCX) fractionation of peptides

Clean peptides were fractionated by 100 mg SCX solid-phase extraction (SPE) columns (HyperSep(tm), Thermo Fisher Scientific) to enhance the peptide spectral information. 600 µg of pooled peptides, including all five types of thyroid tissues (10 N, 42 MNG, 28 FA, 13 FTC, 38 PTC), were reconstituted in equilibration buffer (2.5 mM KH_2_PO_4_ / 25% ACN, pH = 3.0). SCX columns were washed with MilliQ water and equilibration buffer. The pooled sample was then loaded into a conditioned cartridge. Loaded columns were washed with six diluents with different ratios of buffer A (10 mM KH_2_PO_4_ / 25% ACN, pH = 3.0) to buffer B (10 mM KH_2_PO_4_ / 1 M KCl / 25% ACN, pH = 3.0) and increasing KCl concentration. The samples were then split into six fractions and cleaned by C18 spin columns (The Nest Group, United States).

### High-pH reversed-phase chromatography fractionation of peptides

To further increase the peptide overage in the spectral library, another fractionation method, *i*.*e*. high-pH reversed-phase chromatography, was performed. Two pooled samples were combined from 41 follicular thyroid neoplasms (21 FA and 20 FTC) and 16 PTC samples. ∼200 μg of each pooled sample was separated by Thermo Dinex Ultramate 3000 with an XBridge peptide BEH C18 column (4.6 mm X 250 mm, 5 μm, 1 / pkg) at 45 °C. The gradient was 60 min long, with a flow rate of 1 mL/min, and the mobile phase consisting of buffer A (ddH_2_O water with 0.6% ammonia, pH = 10) and buffer B (98% ACN with 0.6% ammonia, pH= 10). The gradient was from 5% to 35% buffer B in condition of pH 10.0 at a flow rate of 1 mL/min. 60 fractions were collected, separated by 1 min interval. The 60 fractions were subsequently combined into 20 fractions for each pooled sample to build the library. The resulting fractionated peptides were then resuspended into 20 µl buffer (2% of ACN, 0.1% formic acid) for instrument injection.

### Data dependent acquisition (DDA)

The fractionated peptides were separated by UltiMate(tm) 3000 RSLCnano System (Thermo Fisher Scientific). The system was equipped with a 15 cm x 75 µm silica column custom packed with 1.9 µm 100 Å C18-Aqua. The mobile phase comprised buffer A (2% ACN, 0.1% formic acid) and buffer B (98% ACN, 0.1% formic acid). Peptides were separated on a 60 min effective liquid chromatography (LC) buffer B gradient (3% to 28% at 300 nL/min). Ionized peptides were transferred into a Q Exactive™ HF MS (Thermo Fisher Scientific). Full MS scans were measured with an Orbitrap at a resolution of 60,000 full widths at half maximum (FWHM) at 200 *m/z* covering 400 to 1200 *m/z* precursors, with automatic gain control (AGC) target value of 3E6 charges and 80 ms maximum injection time (max IT). The top 20 precursor signals were chosen to be fragmented in a higher-energy collision (HCD) cell with 27% normalized collision energy and then transferred to an Orbitrap for MS/MS analysis at a resolution of 30,000 FWHM and an AGC target value of 1E5. By using 60 min LC gradients, we acquired a total of 46 DDA files (Details are listed in Supplementary Table 1).

### Spectral library construction based on DDA

Spectronaut(tm) Pulsar X version 14.6 (Biognosys) was used to generate a spectral library specific to the thyroid. All 46 DDA raw files were searched by Pulsar against a human Swiss-Prot FASTA database (downloaded on 2020-01-22) which included 20,367 protein sequences with FDR of 0.01. The enzyme setting for “trypsin/P” allowed no more than two missed cleavages; cysteine carbamidomethyl was set as a fixed modification, and methionine oxidation was set as a variable modification; mass tolerances were automatically determined, while other settings were left to their default values.

### Quantitative analysis of thyroid samples by data independent acquisition (DIA) and PulseDIA

Together with paired tissues adjacent to the tumor site, three PTC samples were prepared as previously described. Proteomic data for these test thyroid samples was acquired by DIA or PulseDIA, a gas phase fractionation method^20^. For each run, the LC effective gradient was 45 min long, with 3% to 25% buffer B at 0.3 µL/min. MS1 was performed over an *m/z* range of 390-1010 for the DIA, and 390-1210 for the PulseDIA, with a resolution of 60,000 FWHM, an AGC target of 3E6, and a max IT of 80 ms. MS2 was performed with a resolution of 30,000 FWHM, an AGC target of 1E6, and a max IT of 55 ms. For DIA, 24 isolation windows were performed: 20 with 21 *m/z* wide windows, 2 with 41 *m/z* wide windows, and 2 with 61 *m/z* windows. For PulseDIA, five injections with 24 isolation windows per injection were performed^20^. DIA data were analyzed by Spectronaut(tm) version 14.6; all settings were left to their default values.

## Data Records

DDA raw files (Data Citation 1)

Spectral library files in the formats of xlsx, TSV and CSV (Data Citation 2)

DIA raw files (Data Citation 3)

## Technical Validation

### Libraries evaluation

Our thyroid-specific spectral library comprises 925,330 transition groups, 157,548 precursors, 121,960 peptides, 9941 protein groups, and 9826 proteins from proteotypic peptides. An overview is provided in Table 1. Our library is, therefore, more comprehensive than currently published data which include only 2682 proteins^21^.

**Table 1.** Statistics of the thyroid-specific spectral library

|  | **Library** |
| --- | --- |
| **Transition groups** | 925,330 |
| **Peptide precursors** | 157,548 |
| **Peptides** | 121,960 |
| **Protein groups** | 9941 |
| **Proteotypic proteins** | 9826 |

To assess the quality of our spectral library, we first evaluated the composition and distributions of precursors, peptides, and proteins. In our DIA-MS analysis, the precursor mass range cover 400-1200 *m/z*, and approximately 82% of the precursors are between 400-850 *m/z* (Figure 2A). Precursors primarily display two (53%) or three (37%) charges, and their charge distributions are comparable to those of different spectral libraries (Figure 2B)^22^. 82% peptides are 8 to 20 amino acids long, with a median length of 14 amino acids, consistently with the properties of trypsinized peptides (Figure 2C). We next focused on peptide modifications. Oxidation on methionine, the most common modification in our library, was detected in 22,853 peptides, 121,960 of the total peptides. Sample preparation generated 2818 carbamidomethyled peptides at cysteine residues and 2231 N-terminal acetylated ones (Figure 2D). A total of 7634 proteins were detected with at least three proteotypic peptides, and the majority of proteins were found with more than ten (Figure 2E). Additionally, fragments from *y*-ions were more frequently detected than those from *b*-ions due to the collision mode. Our established spectral library achieved comprehensive peptide and protein coverage with high quality.

**Figure 2.**
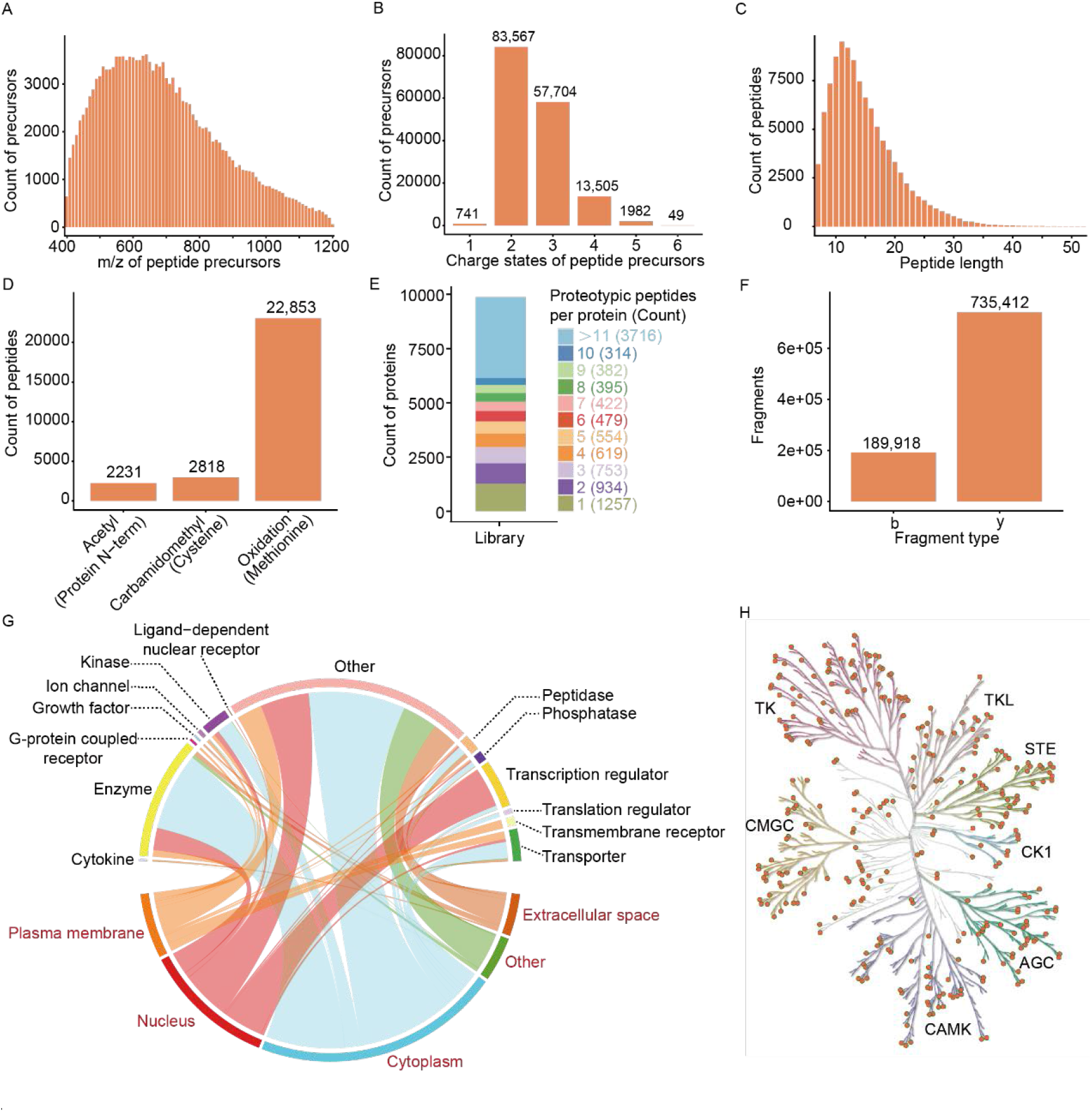
Characterization and statistics of the thyroid-specific spectral library. (A) Distribution of peptide precursor *m/z*. (B) Counts of different precursor charge states. (C) Distribution of identified peptides lengths. (D) Modified peptides numbers and distribution of three modifications. (E) Numbers of proteotypic peptides for each protein and their corresponding ratios and counts. (F) Ion counts of each fragment type. (G) Proteins were annotated according to two classification systems, subcellular location (words in red) and function type (words in black). Each curve represents one protein, linking the protein function type with the corresponding subcellular location. (H) A total of 340 kinases (orange dots), belonging to seven families (highlighted by the different tree colors) were identified in our library.

We next used Gene Ontology to identify the main enriched protein categories within our library. A total of 9,825 proteins were annotated by Ingenuity Pathway Analysis (IPA) software: the enriched protein cellular locations (red words) and protein functions (black words) are shown in Figure 2G. By matching our data to the kinase database KinMap^23^, our library was found to contain 340 kinases from 7 families, accounting for 63.4% (340/536) of the entire kinase database (Figure 2H). These results demonstrate that our library provides a valuable reference for the application of the DIA-MS method to human thyroid samples.

### Technical validation on four datasets

To further validate our library, we analyzed three PTC samples, together with paired tissue samples adjacent to the tumor site. Four datasets were then acquired with the following four acquisition strategies: single-shot DIA (dataset 1), PulseDIA (dataset 2), pre-fraction DIA (dataset 3), and a combination of pre-fraction and PulseDIA (dataset 4). All datasets were subsequently analyzed using Spectronaut 14.6 and our thyroid nodule-specific spectral library. The search results for the four datasets are shown in Figure 3. All three tumor tissues expressed more proteins and peptides than the matched normal thyroid tissues (tumor-adjacent tissues), and this was especially evident at the peptide level (Figure 3A, B). The numbers of identified peptides and proteins using single-shot DIA were the lowest due to the relatively short gradient and the highly abundant protein, thyroglobulin. PulseDIA and pre-fraction DIA led to more identifications. PulseDIA identified more peptides than pre-fraction DIA, but a comparable number of proteins. Finally, a combination of pre-fraction and PulseDIA generated the best results at both peptide and protein levels: 65,544 peptides and 7863 proteins. These results showed that a longer gradient allows the detection of more peptides and proteins.

**Figure 3.**
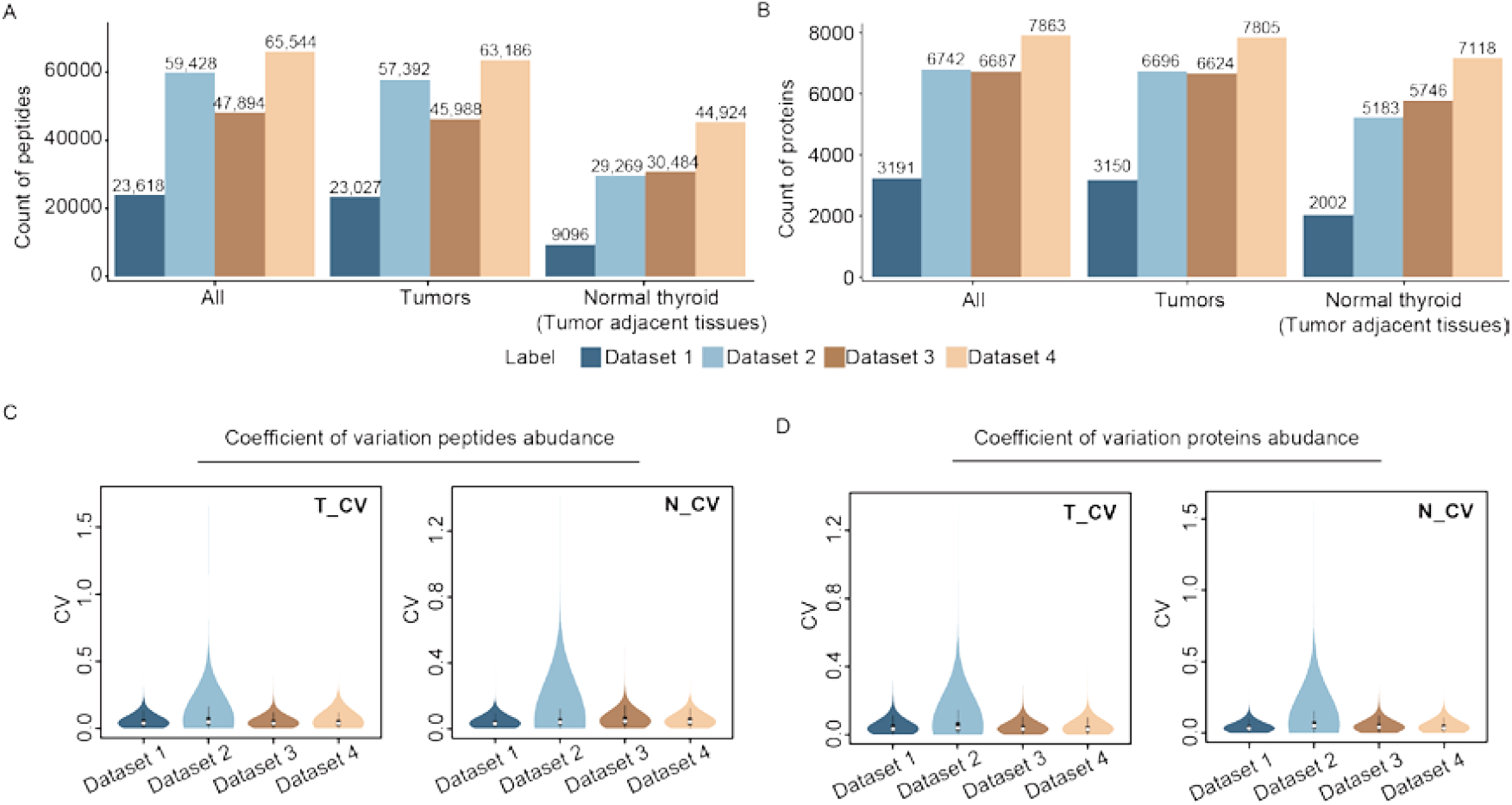
Results from a technical validation of our thyroid-specific spectral library. Four datasets were acquired with single-shot DIA (dataset 1), PulseDIA (dataset 2), pre-fraction DIA (dataset 3), and a combination of pre-fraction and PulseDIA (dataset 4). Identified peptides (A) and proteins (B) obtained by searching against our thyroid specific spectral library. Coefficient of variation of peptides (C) and proteins (D) abundance in tumors (T_CV) and their adjacent normal tissues (N_CV).

We next calculated the coefficient of variation (CV) of peptides and proteins abundance to evaluate the quality of these datasets. The median peptides CVs were less than 0.05 for all datasets (Figure 3C). Similarly, the median proteins CVs were all less than 0.04 (Figure 3D). These results indicate that all four datasets performed well as the quantifications had only negligible differences. These results confirm that our spectral library as a valuable resource provides a robust reference for proteomic exploration of thyroid disease.

Although five types of thyroid tissues and more than 10,000 proteins are in our spectral library, some rare thyroid carcinomas such as anaplastic thyroid carcinoma and medullary thyroid carcinoma were not included in the analysis. This could be addressed in the future with the methodology adopted here. Targeted assays using parallel reaction monitoring (PRM) and selected/multiple reaction monitoring S/MRM could also be developed based on this DIA library^13^. In conclusion, our established DIA library offers a useful resource for proteomic analysis of thyroid tissue specimens.

## Funding

This work is supported by grants from National Key R&0D Program of China (No. 2020YFE0202200), Zhejiang Provincial Natural Science Foundation for Distinguished Young Scholars (LR19C050001), Hangzhou Agriculture and Society Advancement Program (20190101A04), National Natural Science Foundation of China (81972492) and National Science Fund for Young Scholars (21904107). Further support for this project was obtained from the National Cancer Centre Research Fund (Peter Fu Program) and National Medical Research Council Clinician-Scientist Award (NMRC/CSAINV/011/2016).

We thank for the for assistance in data storage, computation and peptide fractionation by the Westlake University Supercomputer Center and the Mass Spectrometry & Metabolomics Core Facility at the Center for Biomedical Research Core Facilities of Westlake University.

## Author contributions

T.G., and Y.S. designed the project. N.G.I. and O.L.K. provided the Singapore set and Y.Z., G.W., Y.H., and J.X. collected the Chinese set. Y.S., L.L., W.L., Q.X and X.C. performed the experiments. Y.S., L.L., W.G. and H.C. conducted proteomic data analysis. Y.S wrote the manuscript, L.L., Z.D., and F.Z. revised the manuscript. T.G. supervised the project.

